## Supplementary Figures and Tables for "Scalable generation of mesenchymal stem cells and adipocytes from human pluripotent stem cells"

##### Figure S1

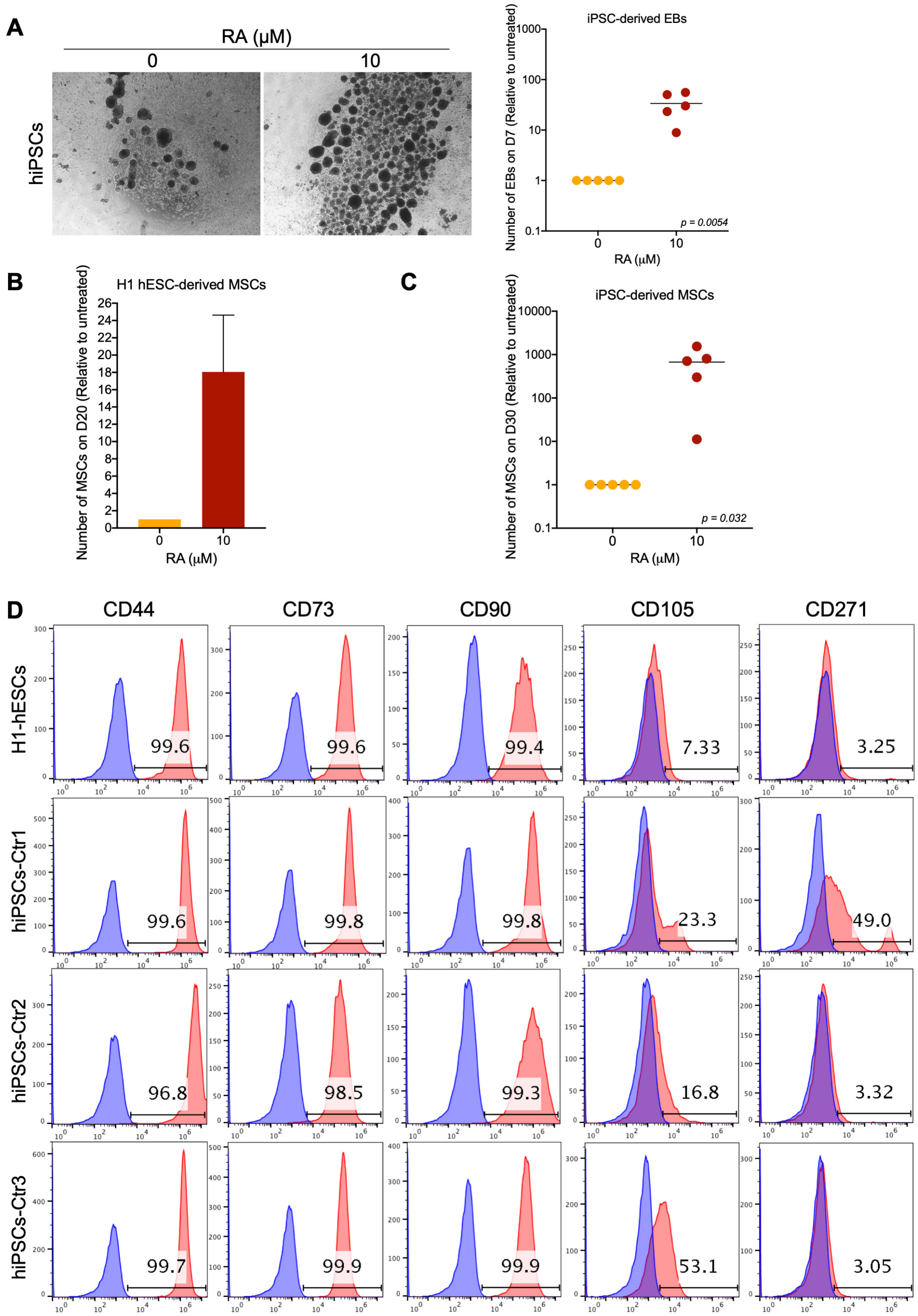

**Supplementary Figure 1.** Effect of short-term treatment with high RA concentration on the efficiency of differentiation of human iPSCs into EBs and MSCs. Human iPSCs (clones generated from three different healthy donors) were differentiated into MSCs as described in Fig 1A of this manuscript without or with treatment with 10 µM RA. A. On day 7 of differentiation, EBs were photographed and counted. Images (10x magnification) for representative hiPSC-derived EBs are shown. The scatter plot presents the individual distributions (dots) and the means (bars) of the fold-increases in EB number (relative to RA untreated condition) on D7 of differentiation of four iPSC clones generated from the PBMCs of three different healthy donors (one to two different clones per donor). B. Scatter plot presenting the fold-increase (relative to RA untreated condition) in the number of MSCs obtained between D25 and D30 of differentiation of three iPSC clones derived from the PBMCs of three healthy donors. Two different representative differentiations were conducted for each clone and presented in the scatter plot. C. Effect of EB treatment with 10 µM RA on the number of MSCs generated from H1-hESCs.The bar graph presents the means ± SD for two independent experiments. D. Flow cytometry analysis of the expression of the MSC markers CD44, CD73, CD90, CD105 and CD271 in the MSCs generated from H1-hESC- and different iPSC-derived EBs treated with 10 µM RA. The histograms presented indicate the percentage of marker-positive cells and are representative for two independent experiments.

**Figure S2**

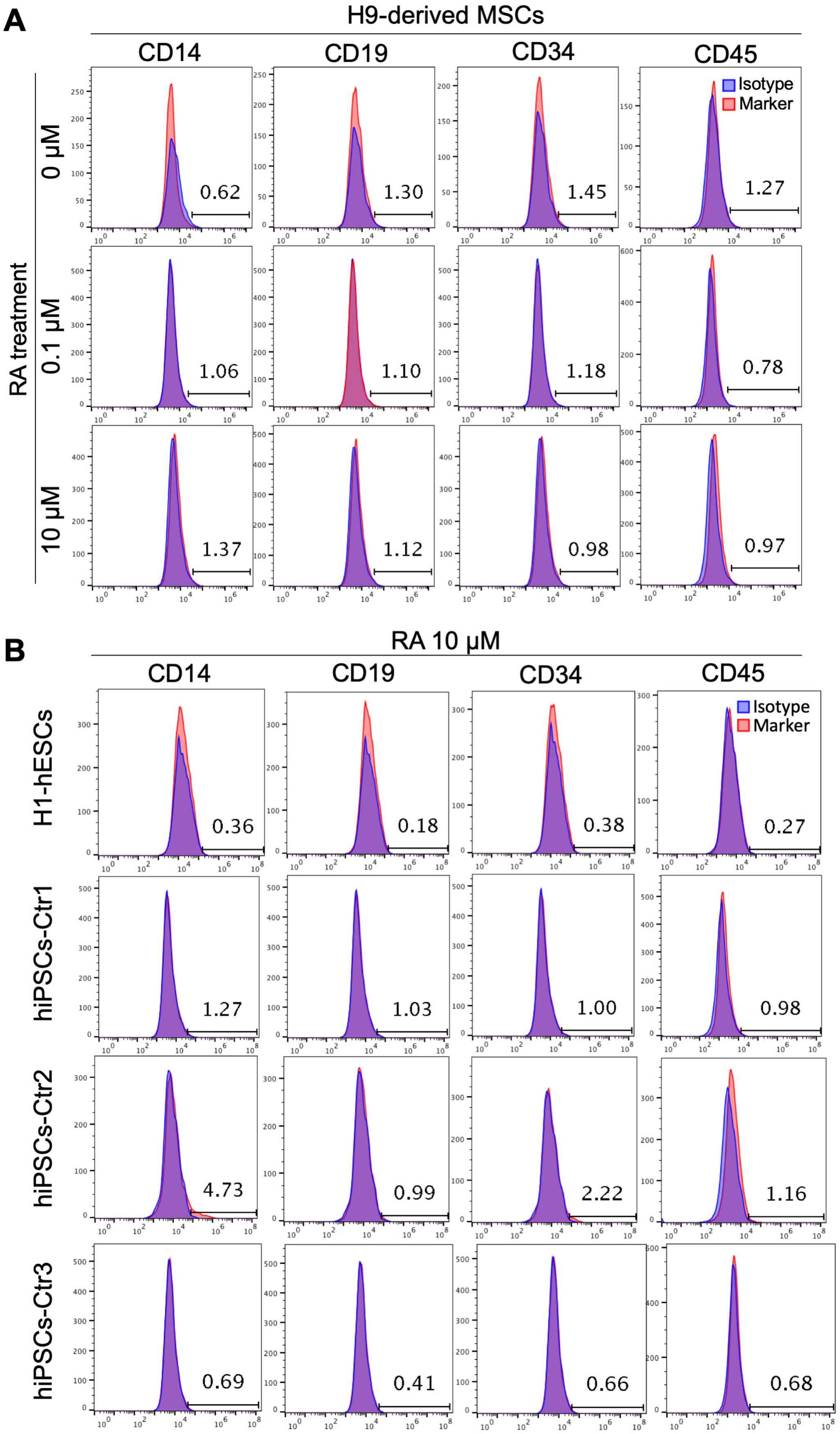
A
H9-derived MSCs
CD14
CD19
CD34
CD45
Isotype
Marker
RA treatment
0 μM
0.1 μM
10 μM
0.62
1.30
1.45
1.27
1.06
1.10
1.18
0.78
1.37
1.12
0.98
0.97
B
RA 10 μM
CD14
CD19
CD34
CD45
Isotype
Marker
H1-hESCs
hiPSCs-Ctr1
hiPSCs-Ctr2
hiPSCs-Ctr3
0.36
0.18
0.38
0.27
1.27
1.03
1.00
0.98
4.73
0.99
2.22
1.16
0.69
0.41
0.66
0.68

**Supplementary Figure 2.** Absence of hematopoietic markers' expression in the PSC-derived MSCs. (A) H9 hESC-derived EBs were treated or not with different RA concentrations and then differentiated into MSCs. The expression of the hematopoietic markers CD14, CD19, CD34 and CD45 was assessed on D20 of differentiation by flow cytometry. The histograms presented indicate the percentage of marker-positive cells and are representative from five-independent experiments. (B) H1 hESC-derived and healthy donor iPSC-derived EBs treated with 10 µM RA and differentiated into MSCs were assessed for the expression of the hematopoietic markers CD14, CD19, CD34 and CD45 on D20 of differentiation by flow cytometry. The histograms presented indicate the percentage of marker-positive cells and are representative from two-independent experiments.

**Fig. S3**

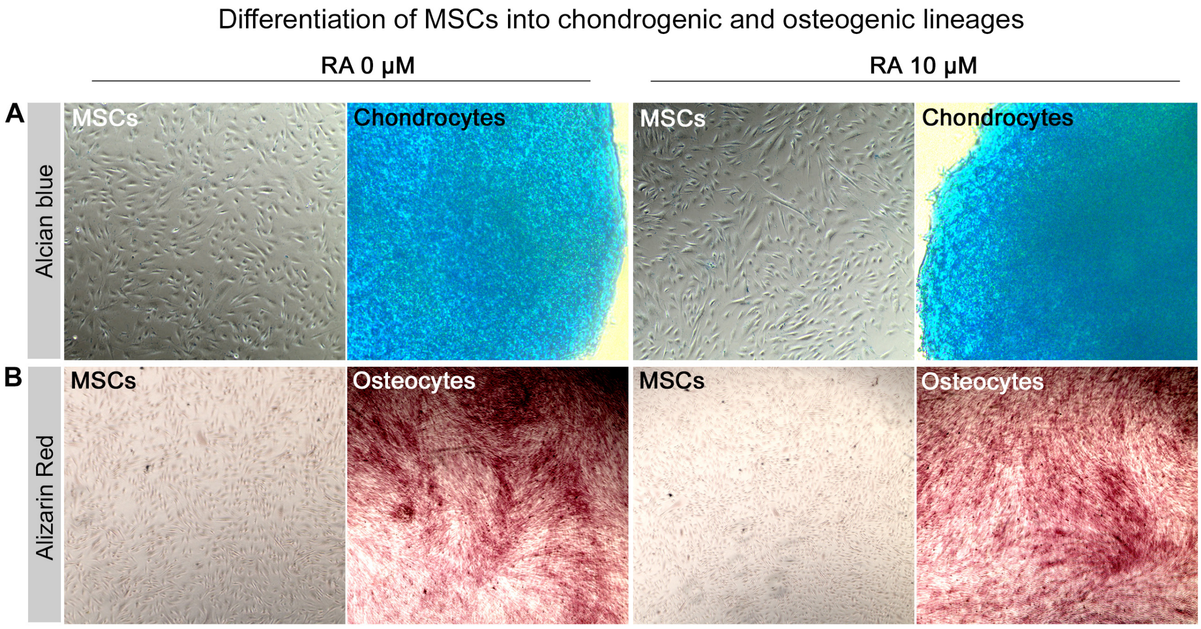


**Supplementary Figure 3.** Differentiation of hPSC-derived MSCs into chondrocytes and osteocytes. MSCs derived from EBs treated or not with 10 µM were differentiated into chondrocytes and osteocytes. Representative images show staining of differentiated cells with alcian blue (A) and alizaran red (B), indicating their successful differentiation into chondrocytes and osteocytes. Undifferentiated MSCs were used as negative control.

**Fig. S4**

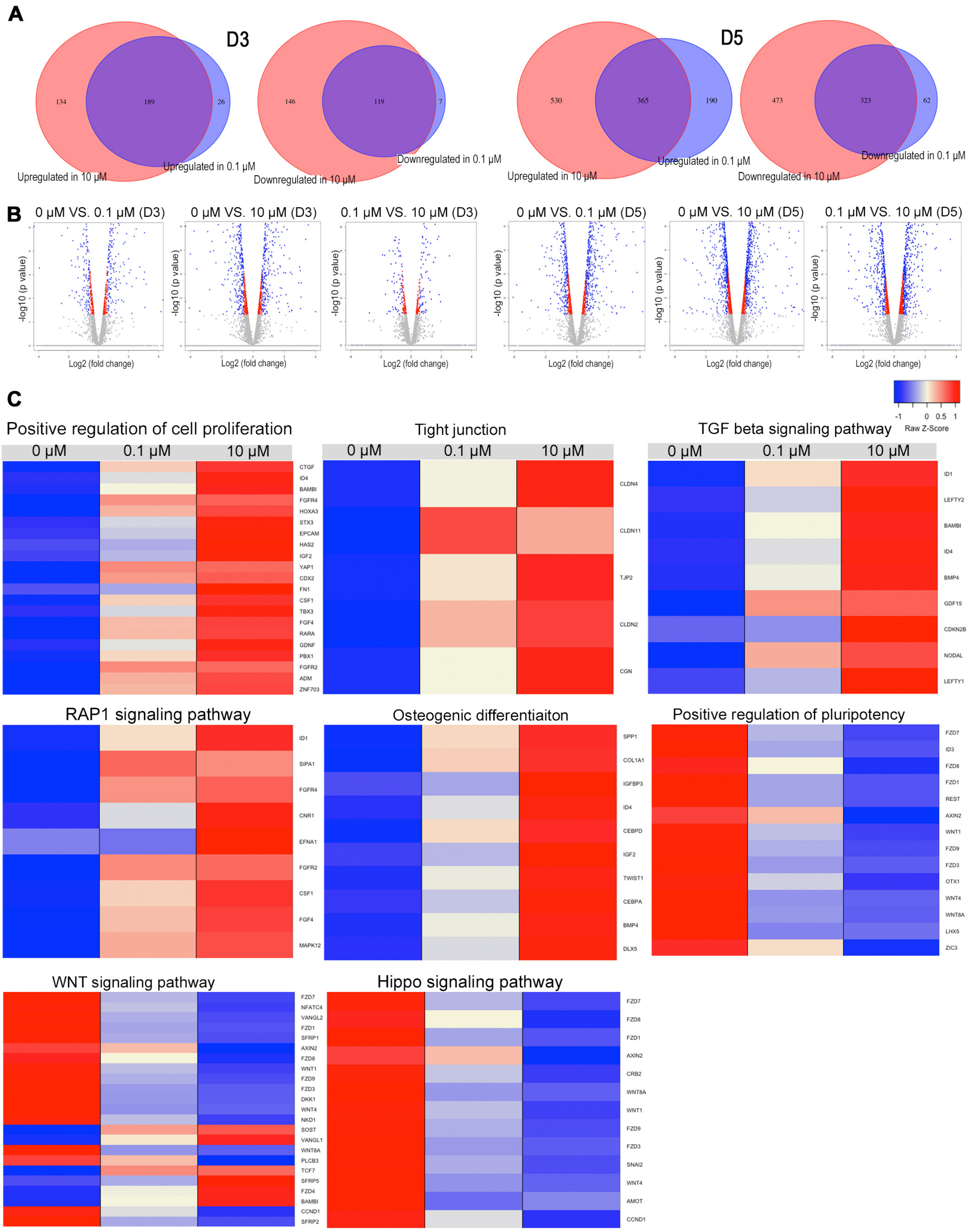


**Supplementary Figure 4.** Transcriptomic comparison of EBs treated with RA versus untreated EBs. (A) Venn diagram showing the number of the DEGs (upregulated and downregulated genes) in H9-hESC-derived EBs treated with 0.1 μM or 10 μM RA in comparison to untreated EBs at day 3 (D3) and day 5 (D5) of differentiation. (B) Volcano plot of the DEGs between RA untreated and RA treated cells at D3 and D5. (C) Heatmaps showing DEGs in H9-hESC-derived EBs treated with 10 μM RA compared to those treated with 0.1 μM RA and untreated EBs at D3 of differentiation. The relative value for each gene is depicted by color intensity, with red indicating upregulated and blue indicating downregulated genes.

### Supplementary Tables

**Table S1. Enriched functions in the upregulated differentially expressed genes (DEGs) in day 5-old EBs treated with 10 μM RA compared with those untreated (*p* <0.05).** n: number of genes from the DEG list in each specific annotation term.

| Gene symbol | Gene title | 10 μM RA vs. 0 μM | |
|---|---|---|---|
| | | Log2 (FC) | p-value |
| **Positive regulation of cell proliferation (n=39)** | | | |
| BNC1 | basonuclin 1 | 4.88 | 5.00E-05 |
| ISL1 | ISL LIM homeobox 1 | 4.71 | 5.00E-05 |
| TBX3 | T-box 3 | 4.21 | 5.00E-05 |
| IGF2 | insulin like growth factor 2 | 4.10 | 5.00E-05 |
| CCKBR | cholecystokinin B receptor | 3.83 | 5.00E-05 |
| HOXA3 | homeobox A3 | 3.67 | 5.00E-05 |
| EPCAM | epithelial cell adhesion molecule | 3.67 | 5.00E-05 |
| KLF5 | Kruppel like factor 5 | 3.39 | 5.00E-05 |
| RAB25 | RAB25, member RAS oncogene family | 2.89 | 5.00E-05 |
| FN1 | fibronectin 1 | 2.82 | 5.00E-05 |
| EDN1 | endothelin 1 | 2.79 | 0.00055 |
| ZNF703 | zinc finger protein 703 | 2.72 | 5.00E-05 |
| NKX3-1 | NK3 homeobox 1 | 2.47 | 0.0026 |
| FGFR4 | fibroblast growth factor receptor 4 | 2.28 | 5.00E-05 |
| TBX2 | T-box 2 | 2.24 | 0.0007 |
| PDGFRA | platelet derived growth factor receptor alpha | 2.12 | 5.00E-05 |
| BAMBI | BMP and activin membrane bound inhibitor | 2.07 | 5.00E-05 |
| SP6 | Sp6 transcription factor | 1.90 | 5.00E-05 |
| HAS2 | hyaluronan synthase 2 | 1.88 | 5.00E-05 |
| PTPN6 | protein tyrosine phosphatase, non-receptor type 6 | 1.78 | 5.00E-05 |
| EGR4 | early growth response 4 | 1.67 | 0.0293 |
| GAB2 | GRB2 associated binding protein 2 | 1.65 | 5.00E-05 |
| CTF1 | cardiotrophin 1 | 1.64 | 0.00045 |
| NCCRP1 | non-specific cytotoxic cell receptor protein 1 homolog | 1.62 | 0.00315 |
| LIF | leukemia inhibitory factor | 1.50 | 0.00155 |
| CLCF1 | cardiotrophin-like cytokine factor 1 | 1.47 | 0.02435 |
| TNC | tenascin C | 1.41 | 5.00E-05 |
| RARA | retinoic acid receptor alpha | 1.37 | 5.00E-05 |
| TGFB1 | transforming growth factor beta 1 | 1.36 | 0.00015 |
| FOSL1 | FOS like 1, AP-1 transcription factor subunit | 1.30 | 0.005 |

| | | | |
|---|---|---|---|
| SOX4 | SRY-box 4 | 1.29 | 5.00E-05 |
| PBX1 | PBX homeobox 1 | 1.29 | 5.00E-05 |
| SPHK2 | sphingosine kinase 2 | 1.23 | 0.03345 |
| STX3 | syntaxin 3 | 1.22 | 5.00E-05 |
| IL6ST | interleukin 6 signal transducer | 1.06 | 0.001 |
| EPHA1 | EPH receptor A1 | 1.04 | 0.0025 |
| CAPN1 | calpain 1 | 0.97 | 0.00165 |
| ITGAV | integrin subunit alpha V | 0.67 | 0.0231 |
| THBS1 | thrombospondin 1 | 0.66 | 0.0302 |
| **Negative regulation of apoptotic process (n=34)** | | | |
| SLC40A1 | solute carrier family 40 member 1 | 6.63 | 0.0232 |
| GATA6 | GATA binding protein 6 | 5.44 | 5.00E-05 |
| OSR1 | odd-skipped related transciption factor 1 | 4.82 | 5.00E-05 |
| TBX3 | T-box 3 | 4.21 | 5.00E-05 |
| KRT18 | keratin 18 | 3.99 | 5.00E-05 |
| TWIST1 | twist family bHLH transcription factor 1 | 3.95 | 5.00E-05 |
| EPCAM | epithelial cell adhesion molecule | 3.67 | 5.00E-05 |
| MMP9 | matrix metallopeptidase 9 | 3.21 | 0.0012 |
| PLK2 | polo like kinase 2 | 3.09 | 5.00E-05 |
| ANXA1 | annexin A1 | 2.89 | 5.00E-05 |
| ID1 | inhibitor of DNA binding 1, HLH protein | 2.52 | 5.00E-05 |
| SMAD6 | SMAD family member 6 | 2.35 | 5.00E-05 |
| DAB2 | DAB2, clathrin adaptor protein | 2.31 | 5.00E-05 |
| SERPINB9 | serpin family B member 9 | 1.91 | 5.00E-05 |
| BMP4 | bone morphogenetic protein 4 | 1.75 | 5.00E-05 |
| RARA | retinoic acid receptor alpha | 1.37 | 5.00E-05 |
| CYR61 | cysteine rich angiogenic inducer 61 | 1.30 | 5.00E-05 |
| EGR3 | early growth response 3 | 1.27 | 0.0037 |
| SPHK2 | sphingosine kinase 2 | 1.23 | 0.03345 |
| NFKBIA | NFKB inhibitor alpha | 1.20 | 0.0008 |
| SOCS3 | suppressor of cytokine signaling 3 | 1.18 | 0.0004 |
| TFAP2A | transcription factor AP-2 alpha | 1.14 | 0.0001 |
| CDKN1A | cyclin dependent kinase inhibitor 1A | 1.08 | 0.0028 |
| GAS6 | growth arrest specific 6 | 1.07 | 0.00355 |
| IL6ST | interleukin 6 signal transducer | 1.06 | 0.001 |
| PRNP | prion protein | 1.06 | 0.0014 |
| SQSTM1 | sequestosome 1 | 1.05 | 0.007 |
| BAG3 | BCL2 associated athanogene 3 | 1.04 | 0.00165 |
| SIAH2 | siah E3 ubiquitin protein ligase 2 | 0.83 | 0.0145 |

| | | | |
|---|---|---|---|
| CD74 | CD74 molecule | 0.82 | 0.0203 |
| PIM1 | Pim-1 proto-oncogene, serine/threonine kinase | 0.81 | 0.00475 |
| VEGFB | vascular endothelial growth factor B | 0.74 | 0.03475 |
| ACTC1 | actin, alpha, cardiac muscle 1 | 0.73 | 0.02365 |
| THBS1 | thrombospondin 1 | 0.66 | 0.0302 |
| **Cell adhesion molecules (CAMs) (n=19)** | | | |
| CLDN2 | claudin 2 | 6.48 | 0.00365 |
| CDH5 | cadherin 5 | 4.93 | 5.00E-05 |
| CLDN4 | claudin 4 | 4.74 | 5.00E-05 |
| CLDN1 | claudin 1 | 3.89 | 5.00E-05 |
| CDH1 | cadherin 1 | 3.19 | 5.00E-05 |
| CDH3 | cadherin 3 | 2.82 | 5.00E-05 |
| CLDN3 | claudin 3 | 2.60 | 5.00E-05 |
| CLDN23 | claudin 23 | 1.91 | 0.00295 |
| SDC4 | syndecan 4 | 1.66 | 5.00E-05 |
| CLDN9 | claudin 9 | 1.59 | 0.02635 |
| PTPRF | protein tyrosine phosphatase, receptor type F | 1.26 | 5.00E-05 |
| HLA-DRB1 | major histocompatibility complex, class II, DR beta 1 | 1.24 | 0.00395 |
| CLDN6 | claudin 6 | 1.17 | 0.00015 |
| HLA-C | major histocompatibility complex, class I, C | 1.11 | 0.0049 |
| F11R | F11 receptor | 1.11 | 0.00015 |
| MPZL1 | myelin protein zero like 1 | 0.98 | 0.00095 |
| NECTIN2 | nectin cell adhesion molecule 2 | 0.79 | 0.00655 |
| ALCAM | activated leukocyte cell adhesion molecule | 0.77 | 0.00895 |
| ITGAV | integrin subunit alpha V | 0.67 | 0.0231 |
| **Tight junction (n=14)** | | | |
| CLDN2 | claudin 2 | 6.48 | 0.00365 |
| CLDN4 | claudin 4 | 4.74 | 5.00E-05 |
| CLDN1 | claudin 1 | 3.88 | 5.00E-05 |
| CRB3 | crumbs 3, cell polarity complex component | 3.76 | 0.01345 |
| CLDN3 | claudin 3 | 2.60 | 5.00E-05 |
| LLGL2 | LLGL2, scribble cell polarity complex component | 2.52 | 5.00E-05 |
| CGN | cingulin | 2.29 | 5.00E-05 |
| CLDN23 | claudin 23 | 1.91 | 0.00295 |
| CLDN9 | claudin 9 | 1.59 | 0.02635 |
| TJP2 | tight junction protein 2 | 1.55 | 5.00E-05 |
| CLDN6 | claudin 6 | 1.17 | 0.00015 |
| F11R | F11 receptor | 1.11 | 0.00015 |
| MYL12B | myosin light chain 12B | 1.10 | 0.00705 |

| | | | |
|---|---|---|---|
| PARD6B | par-6 family cell polarity regulator beta | 0.89 | 0.00965 |
| **ECM-receptor interaction (n=13)** | | | |
| SPP1 | secreted phosphoprotein 1 | 3.45 | 5.00E-05 |
| FN1 | fibronectin 1 | 2.82 | 5.00E-05 |
| ITGA3 | integrin subunit alpha 3 | 2.74 | 5.00E-05 |
| SDC4 | syndecan 4 | 1.66 | 5.00E-05 |
| COL1A1 | collagen type I alpha 1 chain | 1.62 | 5.00E-05 |
| TNC | tenascin C | 1.41 | 5.00E-05 |
| LAMA5 | laminin subunit alpha 5 | 1.37 | 5.00E-05 |
| LAMB1 | laminin subunit beta 1 | 1.28 | 5.00E-05 |
| HSPG2 | heparan sulfate proteoglycan 2 | 0.96 | 0.0052 |
| DAG1 | dystroglycan 1 | 0.78 | 0.00595 |
| ITGAV | integrin subunit alpha V | 0.67 | 0.0231 |
| THBS1 | thrombospondin 1 | 0.66 | 0.0302 |
| SV2A | synaptic vesicle glycoprotein 2A | 0.60 | 0.03965 |
| **Focal adhesion (n=17)** | | | |
| SPP1 | secreted phosphoprotein 1 | 3.45 | 5.00E-05 |
| MYL7 | myosin light chain 7 | 3.19 | 0.02435 |
| FN1 | fibronectin 1 | 2.82 | 5.00E-05 |
| ITGA3 | integrin subunit alpha 3 | 2.74 | 5.00E-05 |
| PDGFRA | platelet derived growth factor receptor alpha | 2.12 | 5.00E-05 |
| COL1A1 | collagen type I alpha 1 chain | 1.62 | 5.00E-05 |
| FLNB | filamin B | 1.47 | 5.00E-05 |
| FLNC | filamin C | 1.45 | 5.00E-05 |
| TNC | tenascin C | 1.41 | 5.00E-05 |
| LAMA5 | laminin subunit alpha 5 | 1.37 | 5.00E-05 |
| LAMB1 | laminin subunit beta 1 | 1.28 | 5.00E-05 |
| MYL12B | myosin light chain 12B | 1.10 | 0.00705 |
| BCAR1 | BCAR1, Cas family scaffolding protein | 0.97 | 0.00145 |
| JUN | Jun proto-oncogene, AP-1 transcription factor subunit | 0.91 | 0.0042 |
| VEGFB | vascular endothelial growth factor B | 0.74 | 0.03475 |
| ITGAV | integrin subunit alpha V | 0.67 | 0.0231 |
| THBS1 | thrombospondin 1 | 0.66 | 0.0302 |
| **Hippo signaling pathway (n=15)** | | | |
| GDF6 | growth differentiation factor 6 | 4.73 | 5.00E-05 |
| WNT6 | Wnt family member 6 | 3.77 | 5.00E-05 |
| FZD4 | frizzled class receptor 4 | 3.36 | 5.00E-05 |
| CDH1 | cadherin 1 | 3.19 | 5.00E-05 |
| LLGL2 | LLGL2, scribble cell polarity complex component | 2.52 | 5.00E-05 |

| | | | |
|---|---|---|---|
| ID1 | inhibitor of DNA binding 1, HLH protein | 1.94 | 5.00E-05 |
| AMOT | angiomotin | 1.75 | 5.00E-05 |
| BMP4 | bone morphogenetic protein 4 | 1.75 | 5.00E-05 |
| TEAD3 | TEA domain transcription factor 3 | 1.55 | 5.00E-05 |
| FZD8 | frizzled class receptor 8 | 1.50 | 5.00E-05 |
| TGFB1 | transforming growth factor beta 1 | 1.36 | 0.00015 |
| FZD5 | frizzled class receptor 5 | 1.17 | 0.0017 |
| SERPINE1 | serpin family E member 1 | 0.94 | 0.0118 |
| PARD6B | par-6 family cell polarity regulator beta | 0.89 | 0.00965 |
| DLG3 | discs large MAGUK scaffold protein 3 | 0.60 | 0.04 |
| **TGF-beta signaling pathway (n=14)** | | | |
| LEFTY2 | left-right determination factor 2 | 9.92 | 5.00E-05 |
| CDKN2B | cyclin dependent kinase inhibitor 2B | 5.30 | 5.00E-05 |
| GDF6 | growth differentiation factor 6 | 4.73 | 5.00E-05 |
| LEFTY1 | left-right determination factor 1 | 4.05 | 5.00E-05 |
| AMHR2 | anti-Mullerian hormone receptor type 2 | 3.34 | 0.001 |
| PITX2 | paired like homeodomain 2 | 3.12 | 5.00E-05 |
| SMAD6 | SMAD family member 6 | 2.35 | 5.00E-05 |
| NODAL | nodal growth differentiation factor | 2.33 | 5.00E-05 |
| BAMBI | BMP and activin membrane bound inhibitor | 2.07 | 5.00E-05 |
| ID1 | inhibitor of DNA binding 1, HLH protein | 1.94 | 5.00E-05 |
| BMP4 | bone morphogenetic protein 4 | 1.75 | 5.00E-05 |
| TGFB1 | transforming growth factor beta 1 | 1.36 | 0.00015 |
| SMAD7 | SMAD family member 7 | 1.12 | 0.00035 |
| THBS1 | thrombospondin 1 | 0.66 | 0.0302 |
| **Positive regulation of epithelial mesenchymal transition (n=7)** | | | |
| TWIST1 | twist family bHLH transcription factor 1 | 3.95 | 5.00E-05 |
| ZNF703 | zinc finger protein 703 | 2.73 | 5.00E-05 |
| DAB2 | DAB2, clathrin adaptor protein | 2.31 | 5.00E-05 |
| BAMBI | BMP and activin membrane bound inhibitor | 2.07 | 5.00E-05 |
| COL1A1 | collagen type I alpha 1 chain | 1.63 | 5.00E-05 |
| TGFB1 | transforming growth factor beta 1 | 1.36 | 0.00015 |
| BCL9L | B-cell CLL/lymphoma 9-like | 0.92 | 0.00155 |

**Table S2. Enriched functions in the downregulated differentially expressed genes (DEGs) in day 5-old EBs treated with 10 μM RA compared with those untreated (*p* <0.05).**
n: number of genes from the DEG list in each specific annotation term.

| Gene symbol | Gene title | 10 μM RA vs. 0 μM | |
|---|---|---|---|
| | | Log2 (FC) | p-value |
| **WNT signaling pathway (n=33)** | | | |
| WNT8A | Wnt family member 8A | -6.74 | 0.02465 |
| FZD10 | frizzled class receptor 10 | -4.11 | 5.00E-05 |
| WNT3A | Wnt family member 3A | -3.16 | 5.00E-05 |
| CCND1 | cyclin D1 | -3.13 | 5.00E-05 |
| NKD1 | naked cuticle homolog 1 | -2.76 | 5.00E-05 |
| ROR1 | Receptor Tyrosine Kinase Like Orphan Receptor 1 | -2.35 | 5.00E-05 |
| LEF1 | lymphoid enhancer binding factor 1 | -2.15 | 5.00E-05 |
| APCDD1 | Adenomatosis Polyposis Coli Down-Regulated 1 Protein | -2.04 | 5.00E-05 |
| LRP4 | LDL Receptor Related Protein 4 | -2.01 | 0.0004 |
| FRZB | Secreted Frizzled-Related Protein 3 | -2.01 | 5.00E-05 |
| WNT5B | Wnt family member 5B(WNT5B) | -1.97 | 5.00E-05 |
| WNT10B | Wnt family member 10B | -1.92 | 5.00E-05 |
| NOTUM | Notum, Palmitoleoyl-Protein Carboxylesterase | -1.92 | 5.00E-05 |
| FZD7 | frizzled class receptor 7 | -1.80 | 5.00E-05 |
| FZD1 | frizzled class receptor 1 | -1.76 | 5.00E-05 |
| GPC4 | glypican 4 | -1.69 | 5.00E-05 |
| FZD3 | frizzled class receptor 3 | -1.68 | 5.00E-05 |
| AXIN2 | axin 2 | -1.61 | 5.00E-05 |
| WNT1 | Wnt family member 1 | -1.51 | 5.00E-05 |
| RSPO3 | R-Spondin 3 | -1.50 | 5.00E-05 |
| TLE4 | TLE Family Member 4, Transcriptional Corepressor | -1.49 | 5.00E-05 |
| NFATC4 | nuclear factor of activated T-cells 4 | -1.33 | 5.00E-05 |
| FZD9 | frizzled class receptor 9 | -1.32 | 0.00775 |
| SFRP2 | secreted frizzled related protein 2 | -1.26 | 5.00E-05 |
| VANGL2 | VANGL planar cell polarity protein 2 | -1.21 | 5.00E-05 |
| SFRP1 | secreted frizzled related protein 1 | -1.03 | 0.00015 |
| DKK1 | dickkopf WNT signaling pathway inhibitor 1 | -0.86 | 0.02085 |
| ROR2 | Receptor Tyrosine Kinase Like Orphan Receptor 2 | -0.85 | 0.0039 |
| MYC | v-myc avian myelocytomatosis viral oncogene homolog | -0.77 | 0.0195 |
| RAC3 | ras-related C3 botulinum toxin substrate 3 | -0.69 | 0.04325 |
| CTNNBIP1 | catenin beta interacting protein 1 | -0.62 | 0.03805 |
| CCND2 | cyclin D2 | -0.61 | 0.02845 |
| LRP6 | LDL receptor related protein 6 | -0.60 | 0.0381 |

| | | | |
|---|---|---|---|
| **Neuronal differentiation and axonal guidance (n=34)** | | | |
| CXCR4 | C-X-C motif chemokine receptor 4 | -4.17 | 5.00E-05 |
| EPHA4 | EPH receptor A4 | -3.21 | 5.00E-05 |
| L1CAM | L1 cell adhesion molecule | -3.07 | 5.00E-05 |
| SHANK1 | SH3 And Multiple Ankyrin Repeat Domains 1 | -2.88 | 5.00E-05 |
| FEZF1 | FEZ Family Zinc Finger 1 | -2.74 | 0.0017 |
| ZIC2 | Zic Family Member 2 | -2.64 | 5.00E-05 |
| ZIC3 | Zic Family Member 3 | -2.57 | 5.00E-05 |
| LRRC4 | leucine rich repeat containing 4 | -2.49 | 5.00E-05 |
| GPM6B | Glycoprotein M6B | -2.37 | 5.00E-05 |
| ZIC1 | Zic Family Member 1 | -2.23 | 5.00E-05 |
| EPHA7 | EPH receptor A7 | -2.16 | 5.00E-05 |
| NGFR | Nerve Growth Factor Receptor | -2.16 | 5.00E-05 |
| ZIC5 | Zic Family Member 5 | -2.16 | 5.00E-05 |
| SEMA5B | semaphorin 5B | -2.10 | 5.00E-05 |
| ASCL1 | Achaete-Scute Family BHLH Transcription Factor 1 | -1.88 | 0.00735 |
| SEMA3G | semaphorin 3G | -1.81 | 0.00035 |
| PAX3 | *Paired Box 3* | -1.74 | 5.00E-05 |
| EFNB1 | Ephrin B1 | -1.65 | 5.00E-05 |
| MDGA1 | MAM Domain Containing Glycosylphosphatidylinositol Anchor 1 | -1.50 | 5.00E-05 |
| SEMA3E | semaphorin 3E | -1.44 | 5.00E-05 |
| NFATC4 | Nuclear Factor Of Activated T Cells 4 | -1.33 | 5.00E-05 |
| SEMA4F | ssemaphorin 4F | -1.09 | 0.00335 |
| SOX11 | SRY-Box Transcription Factor 11 | -1.02 | 0.00035 |
| GAP43 | G protein subunit alpha i2 | -0.97 | 0.0012 |
| KDM7A | Lysine Demethylase 7A | -0.95 | 0.0029 |
| FYN | FYN proto-oncogene, Src family tyrosine kinase | -0.95 | 0.00165 |
| SEMA7A | semaphorin 7A | -0.92 | 0.0071 |
| EFNB3 | Ephrin B3 | -0.90 | 0.0031 |
| DPYSL2 | dihydropyrimidinase like 2 | -0.89 | 0.0028 |
| EFNA3 | ephrin A3 | -0.85 | 0.0201 |
| SEMA3C | semaphorin 3C | -0.85 | 0.00475 |
| TRNP1 | TMF1 Regulated Nuclear Protein 1 | -0.81 | 0.04625 |
| ROBO3 | *Roundabout Guidance Receptor 3* | -0.78 | 0.0431 |
| **Signaling pathways regulating pluripotency of stem cells (n=23)** | | | |
| LHX5 | LIM homeobox 5 | -5.26 | 0.00945 |
| NANOG | Nanog homeobox | -4.17 | 0.00105 |
| WNT8A | Wnt family member 8A | -6.74 | 0.02465 |
| FZD10 | frizzled class receptor 10 | -4.11 | 5.00E-05 |

| | | | |
|---|---|---|---|
| WNT3A | Wnt family member 3A | -3.15 | 5.00E-05 |
| ZIC3 | Zic Family Member 3 | -2.57 | 5.00E-05 |
| SOX2 | SRY-box 2 | -2.15 | 5.00E-05 |
| WNT5B | Wnt family member 5B | -1.97 | 5.00E-05 |
| WNT10B | Wnt family member 10B | -1.92 | 5.00E-05 |
| POU5F1 | POU class 5 homeobox 1 | -1.84 | 5.00E-05 |
| FZD7 | frizzled class receptor 7 | -1.80 | 5.00E-05 |
| FZD1 | frizzled class receptor 1 | -1.76 | 5.00E-05 |
| FZD3 | frizzled class receptor 3 | -1.68 | 5.00E-05 |
| AXIN2 | axin 2 | -1.61 | 5.00E-05 |
| WNT1 | Wnt family member 1 | -1.51 | 5.00E-05 |
| FZD9 | frizzled class receptor 9 | -1.32 | 0.00775 |
| INHBE | inhibin beta E subunit | -1.28 | 0.04045 |
| ACVR2B | activin A receptor type 2B | -0.79 | 0.00515 |
| MYC | v-myc avian myelocytomatosis viral oncogene homolog | -0.77 | 0.0195 |
| FGFR3 | fibroblast growth factor receptor 3 | -0.73 | 0.0152 |
| ID3 | inhibitor of DNA binding 3, HLH protein | -0.71 | 0.0147 |
| RIF1 | replication timing regulatory factor 1 | -0.61 | 0.03265 |
| REST | RE1 silencing transcription factor | -0.60 | 0.03235 |
| **Hippo signaling pathway (n=22)** | | | |
| WNT8A | Wnt family member 8A | -6.73 | 0.02465 |
| SNAI2 | snail family transcriptional repressor 2 | -4.73 | 5.00E-05 |
| FZD10 | frizzled class receptor 10 | -4.11 | 5.00E-05 |
| GDF7 | growth differentiation factor 7 | -3.32 | 5.00E-05 |
| WNT3A | Wnt family member 3A | -3.15 | 5.00E-05 |
| CCND1 | cyclin D1 | -3.13 | 5.00E-05 |
| SOX2 | SRY-box 2 | -2.15 | 5.00E-05 |
| LEF1 | lymphoid enhancer binding factor 1 | -2.14 | 5.00E-05 |
| WNT5B | Wnt family member 5B | -1.97 | 5.00E-05 |
| WNT10B | Wnt family member 10B | -1.92 | 5.00E-05 |
| FZD7 | frizzled class receptor 7 | -1.79 | 5.00E-05 |
| FZD1 | frizzled class receptor 1 | -1.76 | 5.00E-05 |
| FZD3 | frizzled class receptor 3 | -1.68 | 5.00E-05 |
| AXIN2 | axin 2 | -1.60 | 5.00E-05 |
| DLG4 | discs large MAGUK scaffold protein 4 | -1.57 | 0.00885 |
| WNT1 | Wnt family member 1 | -1.51 | 5.00E-05 |
| FZD9 | frizzled class receptor 9 | -1.32 | 0.00775 |
| TGFBR1 | *Transforming Growth Factor Beta Receptor 1* | -0.98 | 0.001 |
| MYC | v-myc avian myelocytomatosis viral oncogene homolog | -0.77 | 0.0195 |

| | | | |
|---|---|---|---|
| TEAD1 | TEA domain transcription factor 1 | -0.67 | 0.01905 |
| GLI2 | GLI family zinc finger 2 | -0.63 | 0.03465 |
| CCND2 | cyclin D2 | -0.61 | 0.02845 |
| **RAP1 signaling pathway (n=14)** | | | |
| FGF3 | fibroblast growth factor 3 | -6.11 | 0.00325 |
| FGF4 | fibroblast growth factor 4 | -4.68 | 0.00325 |
| FGF19 | fibroblast growth factor 19 | -2.67 | 5.00E-05 |
| NGFR | nerve growth factor receptor | -2.16 | 5.00E-05 |
| LPAR4 | lysophosphatidic acid receptor 4 | -2.149 | 5.00E-05 |
| FGF11 | fibroblast growth factor 11 | -1.75 | 5.00E-05 |
| VEGFA | vascular endothelial growth factor A | -1.72 | 5.00E-05 |
| ARAP3 | ArfGAP with RhoGAP domain, ankyrin repeat and PH domain 3 | -1.50 | 0.0002 |
| P2RY1 | purinergic receptor P2Y1 | -1.26 | 0.00055 |
| EFNA3 | ephrin A3 | -0.85 | 0.0201 |
| FGFR3 | fibroblast growth factor receptor 3 | -0.73 | 0.0152 |
| GNAI2 | G protein subunit alpha i2 | -0.71 | 0.0129 |
| RAC3 | ras-related C3 botulinum toxin substrate 3 | -0.69 | 0.04325 |
| GNAS | GNAS complex locus | -0.65 | 0.0203 |

**Table S3: The details of the antibodies used for flow cytometry and immunostaining**

| For Flow Cytometry | | | |
|---|---|---|---|
| **Antibody** | **Company** | **Catalog #** | **Dilution or Concentration** |
| APC anti-mouse/human CD44 | Biolegend | 103012 | 1:100 |
| APC anti-human CD73 (Ecto-5'-nucleotidase) | Biolegend | 344006 | 1:100 |
| APC anti-human CD90 (Thy1) | Biolegend | 328114 | 1:100 |
| APC anti-human CD105 | Biolegend | 323208 | 1:100 |
| APC anti-human CD271 (NGFR) | Biolegend | 345108 | 1:100 |
| PE anti-human CD45 | Biolegend | 304008 | 1:100 |
| FITC anti-human CD14 | Biolegend | 301804 | 1:100 |
| FITC anti-human CD19 | Biolegend | 302206 | 1:100 |
| FITC anti-human CD34 | Biolegend | 343504 | 1:100 |
| Mouse anti-FABP4 | Abcam | ab93945 | 1:100 |
| Rabbit anti-Sox2 | ThermoFisher Scientific | 48-1400 | 1:100 |
| Alexa Fluor 647 anti-mouse IgG | ThermoFisher Scientific | A31571 | 1:500 |
| Alexa Fluor 488 anti-rabbit IgG | ThermoFisher Scientific | A21206 | 1:500 |
| Mouse IgG1, κ Isotype Ctrl | Abcam | ab91353 | 1:100 |
| APC Mouse IgG1, κ Isotype Ctrl | Biolegend | 400122 | 1:100 |
| APC Rat IgG2b, κ Isotype Ctrl | Biolegend | 400612 | 1:100 |
| PE Mouse IgG1, κ Isotype Ctrl | Biolegend | 400114 | 1:100 |
| FITC Mouse IgG2a, κ Isotype Ctrl | Biolegend | 400208 | 1:100 |
| FITC Mouse IgG1, κ Isotype Ctrl | Biolegend | 400107 | 1:100 |
| **For immunostaining** | | | |
| Rabbit anti-PPARγ | CST | 2443 | 1:1000 |
| Goat anti-FABP4 | R & D Systems | AF3150 | 1:200 |
| Mouse anti-adiponectin | Abcam | ab22554 | 1:400 |
| Alexa Fluor 488 anti-goat IgG | ThermoFisher Scientific | | 1:500 |
| Alexa Fluor 568 anti-rabbit IgG | ThermoFisher Scientific | A-10042 | 1:500 |
| Alexa Fluor 568 anti-mouse IgG | ThermoFisher Scientific | A-10037 | 1:500 |
